# Cardiomyocyte ploidy is dynamic during postnatal development and varies across genetic backgrounds

**DOI:** 10.1101/2022.09.15.508152

**Authors:** Samantha K. Swift, Alexandra L. Purdy, Mary E. Kolell, Michael A. Flinn, Caitlin Lahue, Tyler Buddell, Kaelin A. Akins, Parker Foster, Caitlin C. O’Meara, Christoph D. Rau, Michaela Patterson

## Abstract

Somatic polyploidization, an adaptation by which cells increase their DNA content to support cell and organ growth, is observed in many mammalian cell types, including cardiomyocytes. Although polyploidization is beneficial in many contexts, progression to a polyploid state is often accompanied by a loss of proliferative capacity. Recent work suggests that heterogeneity in cardiomyocyte ploidy is highly influenced by genetic diversity. However, the developmental course by which cardiomyocytes reach their final ploidy state has only been investigated in select genetic backgrounds. Here, we assessed cardiomyocyte number, cell cycle activity, and ploidy dynamics across two divergent inbred mouse strains; C57Bl/6J and A/J. Both strains are born and reach adulthood with a comparable number of cardiomyocytes, however the end composition of ploidy classes and developmental progression to reach the final state and number differ substantially. In addition to corroborating previous findings that identified *Tnni3k* as a mediator of cardiomyocyte ploidy, we also uncover a novel role for *Runx1* and *Tnni3k* in ploidy dynamics and cardiomyocyte cytokinesis. These data provide novel insight into the developmental path to cardiomyocyte ploidy states and challenge the paradigm that polyploidization and hypertrophy are the only mechanisms for growth in the mouse heart after the first week of life.

## INTRODUCTION

Somatic polyploidization, a cellular process resulting in the retention of multiple copies of the archetypical diploid genome, is a key component of development and organogenesis for many mammalian tissues, including the heart. Cardiomyocytes transition to a polyploid state beginning around birth, with the exact timing being species specific (Gan et al., 2020; Patterson and Swift, 2019). Polyploidization largely coincides with the shift from hyperplastic to hypertrophic growth of the myocardium (Soonpaa et al., 1996). While the exact function of somatic polyploidy in cardiomyocytes is still not fully understood, it has been implicated in energy preservation during rapid postnatal growth, maintenance of intercalated discs and the pseudosyncitium, establishment of greater force-generating muscle units, and terminal maturation (Orr-Weaver, 2015; Patterson and Swift, 2019). Recent literature, links cardiomyocyte polyploidization with loss of myocardial regenerative competence (Gonzalez-Rosa et al., 2018; Han et al., 2020; Patterson et al., 2017). Insights into the developmental progression to cardiomyocyte polyploidy would improve our understanding of specific aspects of myocardial biology including total cardiomyocyte number, cell cycle potential in naïve and disease contexts, and capacity for myocardial regeneration.

Cardiomyocyte polyploidy arises via an alternative cell cycle known as endomitosis, in which cells replicate their genome without completing mitosis. Insights from mouse studies suggest cardiomyocyte polyploidization is tightly linked to cell cycle exit. For example, in the mouse, cardiomyocyte completion of cytokinesis rapidly declines within the first 1-2 days post birth and DNA synthesis during the first postnatal week largely contributes to cardiomyocyte polyploidization. Subsequently, additional DNA synthesis ceases around postnatal day (P) 10 at which point both the ploidy state of individual cardiomyocytes and the final number of cardiomyocytes are thought to be largely determined and constant (Alkass et al., 2015; Soonpaa *et al*., 1996; Soonpaa et al., 2015; Walsh et al., 2010). The timeline of cell cycle exit is further supported by the loss of cardiac regenerative capacity after P7 in mice (Porrello et al., 2011). Strikingly, ploidy in other cell types, such as hepatocytes, is not believed to be static as has been proposed in cardiomyocytes, but instead a fluid and dynamic state (Duncan, 2013).

Cardiomyocytes display diverse ploidy states. A single round of endomitosis results in cells with twice as much DNA (i.e. 4N), while a second round of endomitosis would produce 8N cells. Another layer of complexity arises from the stage at which a cardiomyocyte exits the cell cycle. Cardiomyocytes can exit the endomitotic cell cycle just prior to karyokinesis resulting in single nucleus, polyploid cells (1×4N, 1×8N, etc.). Conversely cardiomyocytes that successfully complete karyokinesis but fail to complete cytokinesis result in multinucleated cells with diploid nuclei (2×2N, 4×2N, etc.). In some cases, a combination of the two exit points ensues (2×4N, or Trinucleated 1×4N;2×2N). Together, these cell fate decisions contribute to a final composition of diverse ploidy classes, which display both inter- and intraspecies variation. For example, human cardiomyocytes are predominantly mononuclear and polyploid (Mollova et al., 2013) while murine cardiomyocytes are predominately binucleated (Soonpaa *et al*., 1996) and porcine cardiomyocytes can have upwards of 16 diploid nuclei (Velayutham et al., 2020). An interesting question that has arisen from the field is if the various ploidy classes bestow distinct attributes to the myocardium. Recent literature suggests that having a higher proportion of mononuclear diploid cardiomyocytes (MNDCMs, 1×2N) is associated with greater regenerative competence, while polyploidy in cardiomyocytes impairs the proliferative response (Gonzalez-Rosa *et al*., 2018; Han *et al*., 2020; Hirose et al., 2019; Patterson *et al*., 2017). Beyond regeneration, the roles distinct ploidy classes play in cardiac homeostasis and pathophysiology remains largely unexplored.

Much of our understanding surrounding the timing and progression of polyploidy stems from work on mice and is limited to only a few strains. We recently determined that ploidy class ratios vary dramatically across inbred mouse strains, where some strains have a higher proportion of MNDCMs, while other strains display higher proportions of cardiomyocytes with ≥8N DNA content (Patterson *et al*., 2017). These findings suggest that genetics influence cardiomyocyte ploidy composition and raise the concern that our insights into polyploid progression and cardiomyocyte cell cycle dynamics may be hampered by only examining the process in select strains. We initiated the experiments described here hypothesizing that two strains of mice with divergent ploidy composition in adulthood arise at their terminal states via distinct approaches.

## RESULTS

### Polyploidization of C57Bl/6J and A/J cardiomyocytes follow distinct developmental programs

We sought to characterize the progression of cardiomyocyte polyploidy from the early postnatal period through adulthood across two genetically and phenotypically divergent, inbred mouse strains, A/J and C57Bl/6J. These strains were selected based on Patterson et. al.’s demonstration that A/J had 5-fold higher MNDCM content than C57Bl/6J at 6 weeks of age. First, we established the total number of cardiomyocytes in both C57Bl/6J and A/J ventricles at multiple time points, ranging from postnatal day 1 (P1) to 6 weeks of age (Figure 1A, and Supp Table 1). Cardiomyocytes were counted via hemocytometer and distinguished from non-cardiomyocytes by morphology and size. We found C57Bl/6J mice had a significant increase in the number of cardiomyocytes from P1 to P7 (P < 0.0001), after which total cardiomyocyte number displayed minimal expansion (P = 0.077 P7 to 6 week). This result is consistent with previous literature for C57Bl/6 mice (Alkass *et al*., 2015; Soonpaa *et al*., 2015) and suggests that some residual completion of the canonical mitotic cell cycle takes place after birth. Conversely, A/J ventricles demonstrated a slower, though still significant, initial increase in cardiomyocyte numbers in the week immediately following birth (P = 0.041). Unlike C57Bl/6J mice, a second significant increase in cardiomyocyte numbers occurred from P21 to 6 weeks in A/J mice (P = 0.036) (Figure 1A).

**Figure 1.**
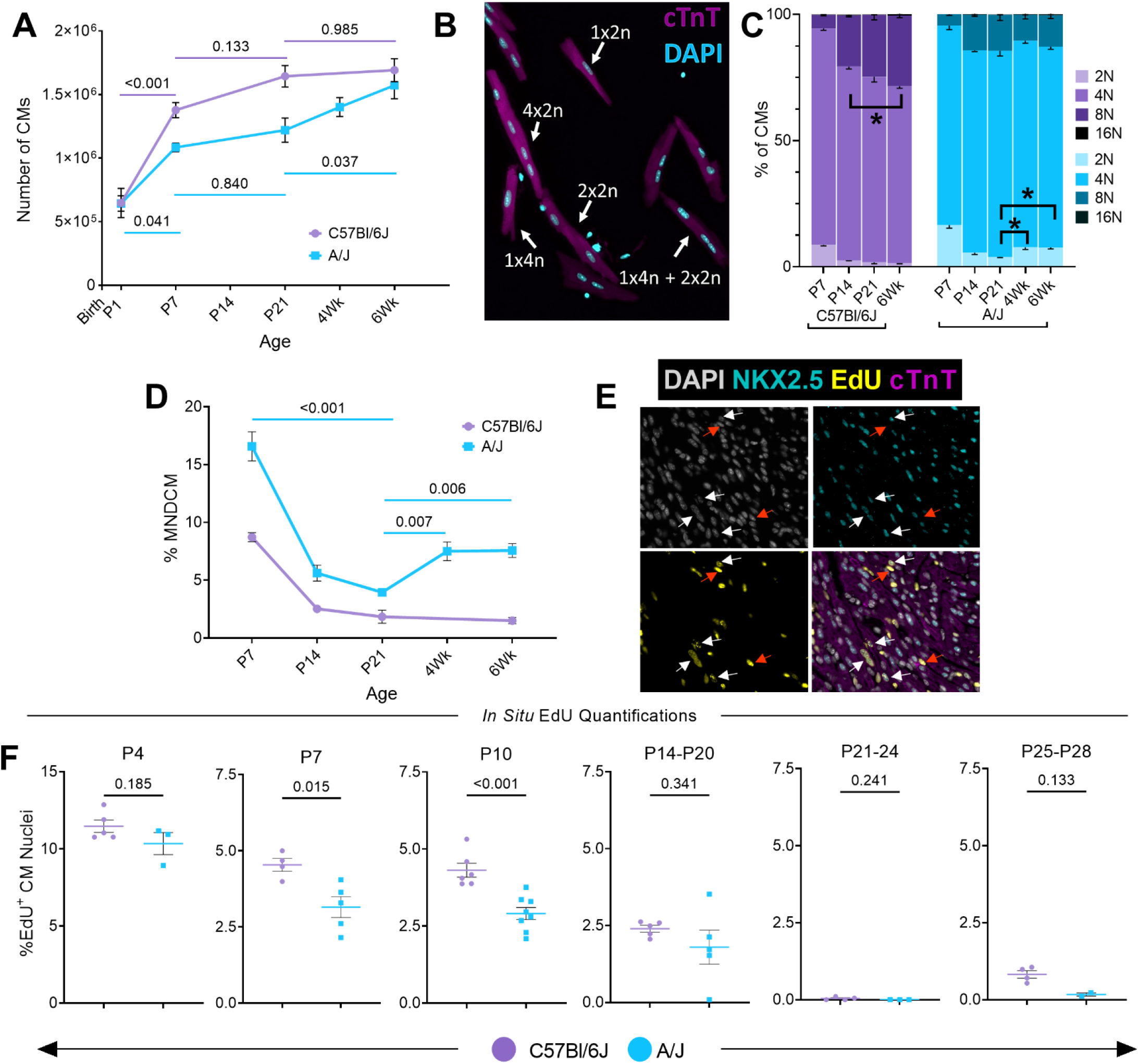
Developmental progression of polyploidy across to genetically divergent mouse strains. **(A)** Number of cardiomyocytes counted via hemocytometer after Langendorff dissociation over multiple timepoints from P1 to 6 weeks in C57Bl/6J (purple) and A/J mice (blue) (N=6-14). Complete breakdown of data and statistical comparisons can be found in Supp Table 1. **(B)** Single-cell ventricular suspension stained for cardiac troponin T (cTnT) (magenta) and DAPI (cyan). Identified cardiomyocyte ploidy classes labeled on the image. **(C)** Quantification of 2N (1×2N), 4N (sum of 1×4N and 2×2N populations), 8N (sum of 1×8N, 2×4N, Tri: 1×4N + 2×2N, and 4×2N populations), and 16N (sum of 1×16N, 2×8N, Tri: 1×8N + 2×4N, and 4×4N populations) ploidy classes at multiple timepoints from P7 to 6 weeks in C57Bl/6J and A/J mice (N=3-9). * indicates p<0.05 for select comparisons; complete statistical comparisons can be found in Supp Table 2. **(D)** Percent mononuclear diploid cardiomyocytes (MNDCMs) extracted from Figure 1C (N=3-9). **(E)** Representative image of cardiomyocyte DNA synthesis measured *in situ* with DAPI (grey), NKX2.5 (cyan), EdU (yellow), and cTnT (magenta). White arrows indicated EdU, NKX2.5-double positive cardiomyocyte nuclei, red arrows are EdU-positive NKX2.5-negative non-cardiomyocyte nuclei. **(F)** Quantification of the percentage of EdU, NKX2.5-double positive nuclei as a percent of total NKX2.5-positive cardiomyocyte nuclei in the left ventricle in both strains across multiple timepoints. Timepoints above each graph indicate the time of the EdU injection. Tissue was collected 24 hours after last injection.

Alongside our assessment of total cardiomyocyte number, we examined the dynamics of polyploidization across the two strains from P7 through 6 weeks of age. In single cell suspensions generated from Langendorff digested ventricles we quantified various ploidy classes, including MNDCMs (2N – 1×2N), tetraploid cardiomyocytes (4N – 1×4N and 2×2N), octoploid cardiomyocytes (8N – 1×8N, 2×4N, trinucleated – 1×4N + 2×2N, and tetranucleated – 4×2N), and a rare 16N cardiomyocyte (Figure 1B-C). This analysis revealed that C57Bl/6J mice display a substantial increase in cardiomyocyte polyploidization from P7 to P14 at which point only 2.5% of cardiomyocytes remained mononuclear and diploid (Figure 1C-D). The vast majority of cardiomyocytes are 4N by P7 and this ploidy class remains the majority at 6 weeks. Octoploid cardiomyocytes most strikingly increase in number during the second postnatal week (from P7 to P14), suggesting that a second round of endomitosis takes place during this time period. Very little change in ploidy states occurs after P14 beyond a gradual expansion of the 8N population from P14 to 6 weeks (P=0.037) (Figure 1C, Supp Table 2). Meanwhile, A/J cardiomyocyte ploidy increased until P21 when frequency of a residual MNDCM population reached its lowest point and the frequency of the 8N population its maximum (Figure 1C-D). Surprisingly, between P21 and 6 weeks of age the MNDCM population nearly doubled in size from 3.9% at P21 to 7.6% at 6 weeks (P=0.009), a phenomenon which was not observed in C57Bl/6J (Figure 1D). To narrow the window of when this expansion occurred, we added a 4-week-old collection timepoint with A/J mice, demonstrating that the increase of the MNDCM population largely took place during the 4^th^ postnatal week of life. This expansion coincided with the second wave of increased cardiomyocyte numbers unique to A/J mice (Figure 1A). These observed cellular differences between strains did not impact heart weight nor heart-weight-to-body-weight ratios at the time points assessed (Supp Figure 1).

To identify an explanation for the robust expansion of MNDCMs after P21 in A/J mice, we assessed DNA synthesis via intraperitoneal (i.p.) injections of EdU at select timepoints throughout postnatal development and evaluated by *in situ* immunofluorescence (Figure 1E) one day following the final injection (Figure 1F). Both C57Bl/6J and A/J hearts displayed the highest level of DNA synthesis at P4. DNA synthesis reached near negligible levels in both strains by the 3^rd^ postnatal week, consistent with past reports (Soonpaa *et al*., 1996; Soonpaa *et al*., 2015). At P7 and P10, A/Js showed significantly reduced numbers of EdU-positive cardiomyocytes. This aligned with our observation of a slower increase in total cardiomyocyte number compared to C57Bl6Js (Figure 1A) and slower conversion to polyploid states (Figure 1C-D). When labeling in multiple-day increments from P21 onward, very few EdU-positive cardiomyocytes were identified in either strain (Figure 1F). Further, we detected no increase in DNA synthesis in A/J compared to C57Lb/6J during the 3–4-week timepoint, suggesting the robust expansion of A/J cardiomyocytes observed between P21 and 4 weeks could not be explained by traditional mitotic expansion of the MNDCM population.

### A/J cardiomyocytes display ploidy reversal following weaning

To investigate the expansion of MNDCM frequency in A/J mice between P21 and 6 weeks of age in the absence of new DNA synthesis, we quantified the completion of cytokinesis by a single cell suspension method in both A/J and C57Bl/6J mice (Figure 2A-C). Briefly, if a cardiomyocyte labeled with EdU is both mononuclear and diploid it is interpreted to have completed cytokinesis (Figure 2A, (Auchampach et al., 2022)). We used this logic to see if we could detect 1×2N EdU-positive cardiomyocytes at a timepoint after the observed MNDCM expansion at 6 weeks, which were not present at P21. To achieve this, we labeled cardiomyocytes with daily 10mg/kg injections of EdU from P14-20. Following EdU administration, we analyzed nucleation and ploidy of EdU-positive cardiomyocytes at two timepoints, P21 or 6 weeks of age by single cell suspension methods (Figure 2B-C). With this EdU regimen, P21 A/J ventricles displayed a slight reduction in the frequency of EdU-positive cardiomyocytes compared to C57Bl/6J ventricles (Figure 2D). This decreased EdU incorporation was not observed by *in situ* quantification methods with a similar injection strategy but is consistent with observations at other timepoints assessed *in situ* (i.e. P7 and P10, Figure 1F). In line with our previous data, and the current literature, C57Bl/6J mice had little to no EdU-positive MNDCMs at either the P21 or 6-week collection timepoint (Figure 2E), suggesting EdU injections in the 3^rd^ postnatal week do not contribute to cell division. Instead, the vast majority of C57Bl/6J cardiomyocytes undergoing DNA synthesis during the 3^rd^ postnatal week are becoming 8N (Figure 2F), likely arising from a 4N cell undergoing a second round of endomitosis. In contrast, Edu-positive MNDCMs can be found at both P21 and 6 weeks of age in A/J mice, suggesting some completion of cell division is still taking place. Most surprisingly, despite identical EdU injection regimens and random segregation of littermates across timepoints to avoid batch effects, we quantified a more than two-fold increase in EdU+ MNDCMs at the 6-week timepoint compared to P21 in A/J hearts (~2.5% to ~6%, P=0.04, Figure 2E). These data imply that a portion of cardiomyocytes which underwent DNA synthesis during the 3^rd^ postnatal week is not completing cytokinesis until after P21.

**Figure 2.**
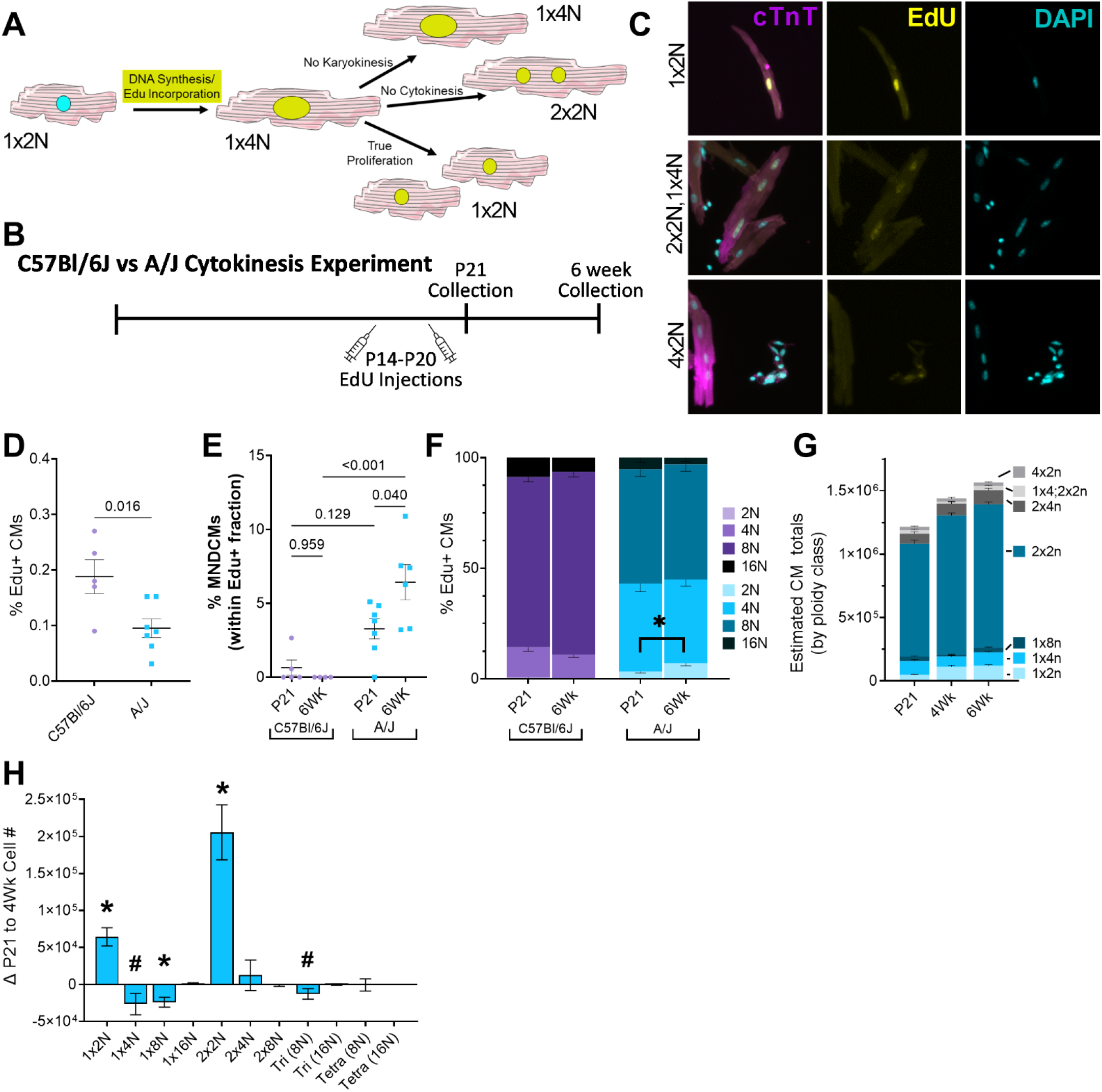
A/J cardiomyocytes undergo a cytokinetic event after P21 resulting in expansion of the MNDCM population. **(A)** Schematic depicting possible outcomes after EdU labeling. EdU-positive cardiomyocytes can only become mononuclear and diploid if cytokinesis was successfully completed. **(B)** Timeline of EdU injections and cell collection timepoints. **(C)** Representative images of single cell ventricular suspensions stained for cTnT (magenta), EdU (yellow), and DAPI (cyan). Tri- and tetranucleated cardiomyocytes are used to normalize DAPI fluorescence intensity. Top: EdU-positive MNDCM (1×2N); middle: EdU-positive trinucleated cardiomyocyte (2×2N; 1×4N); and bottom: EdU-negative tetranucleated cardiomyocyte (4×2N). **(D)** Quantification of total EdU-positive cardiomyocytes following EdU administration outlined in Figure 2B represented as a percent of total cardiomyocytes. Assessed at P21 collection timepoint. **(E)** Quantifications of EdU-positive MNDCMs as a percent of total EdU-postive cardiomyocytes across both A/J and C57Bl/6J at P21 and 6 weeks. **(F)** Quantifications of EdU+ cardiomyocytes broken down into total DNA content (i.e. 2N, 4N, 8N, or 16N). **(G)** Estimated total number of cardiomyocytes in A/J for each ploidy class. Calculations were determined by multiplying the total number of cardiomyocytes (Figure 1A) by the percentage of each ploidy class (Figure 1C) at P21, 4, and 6 weeks. **(H)** Delta change between P21 and 4 weeks in cardiomyocyte number within each ploidy class. * indicates P<0.05; # indicates P≤0.1.

To refine which weeks of postnatal development were contributing to cardiomyocyte cytokinesis in A/Js, we conducted two additional cytokinesis experiments in A/J alone, labeling with EdU during earlier developmental timepoints either at the first (Experiment [Exp] #1) or the second (Exp #2) postnatal weeks, respectively, and assessing for cytokinesis at either P21 or 4 weeks (Supp Figure 2A). When labeling in the first week at P4 and P5, a similar phenotype was observed as with the third postnatal week labeling strategy: EdU-positive MNDCMs were more frequently identified at four weeks (3.03%) compared to P21 (1.18%) (Supp Figure 2B, P=0.001). However, there were nominal numbers (<1%) of EdU-positive MNDCMs identified at P21 and 4 weeks when labeling in the second postnatal week (Supp Figure 2D), indicating the DNA synthesis occurring during the second postnatal week primarily contributes to polyploidization of cardiomyocytes. More specifically, 80% of cardiomyocytes undergoing DNA synthesis in this second postnatal week are becoming ≥8N (Supp Figure 2E). This was a greater percentage of ≥8N CMs than was seen with injection regimens at any other time point (Figure 2F, Supp Figure 2C).

Upon confirmation of cytokinesis in A/J mice, we attempted to elucidate which ploidy classes were most highly contributing to the expansion of total cardiomyocytes and the MNDCM ploidy class. To derive an estimate of the total number of cardiomyocytes in each ploidy class at each time point, the ploidy class percentages at P21, 4 and 6 weeks were multiplied by the total cardiomyocyte numbers at each respective time point (Figure 2G). With this calculation, we could again confirm that the 1×2N population had expanded from P21 to 4 weeks of age by a mean of ~64,000 CMs (Figure 2G-H, P<0.0002). We also detected a significant increase in 2×2N population. As for populations that were decreasing over this same period, only the 1×8N population reached statistical significance decreasing by ~24,000 cells between P21 and 4 weeks (P=0.002), however other populations also decreased with less confidence, including the 1×4N (~26,000; P=0.10) and trinucleated, 1×4N + 2×2N populations (~12, 000; P=0.08) (Figure 2H). These results suggest that 1×8N cardiomyocytes, and possibly additional polyploid populations, are contributing to the expansion of the MNDCM population, a possibility which would infer that “ploidy reversal” (Duncan et al., 2010) is taking place.

Past analysis from just one or two mouse strains, including C57Bl/6, has suggested that cardiomyocyte ploidy is largely static after about P14. Our detailed analysis presented here suggest a much more dynamic process in A/J hearts, whereby a subset of cardiomyocytes which undergo DNA synthesis in the first postnatal week and again in the third postnatal week, complete cytokinesis sometime after weaning, a possibility which would be indicative of the ploidy conveyor model put forth by the hepatocyte field. Mathematically, this could add up as well. If an 8N cell is capable of reverting to a 2N state through multipolar spindles (Duncan *et al*., 2010), it would generate up to 4 new daughter cells in the process. Therefore, our ~24,000 1×8N cells would become 96,000 1×2N. It remains possible that other reversion combinations also ensue. For example, perhaps some of the 8N cells only revert to a 4N state this would explain why we see such a prominent 4N population in the single cell EdU analysis (Figure 2F, and Supp Figures 2C and 2E).

### Tnni3k hypomorphism is partially responsible for ploidy dynamics

We next began to explore possible mechanisms for this unique and dynamic ploidy phenotype observed in A/J mice. A/J mice harbor a naturally occurring hypomorphic mutation in the gene *Tnni3k (Wheeler et al., 2009*), while C57Bl/6J carry the “wild-type” variant. Utilizing an engineered knockout allele of *Tnni3k* maintained on a C57Bl/6J inbred background, Patterson et al. (2017) previously established that genetic ablation of *Tnni3k* contributes to MNDCM frequency. Specifically, *Tnni3k* knockout mice displayed a MNDCM frequency of 5.3% in early adulthood compared to just 1.5% in controls (Patterson *et al*., 2017). Additional loss-of-function alleles of *Tnni3k* have yielded similar results (Gan et al., 2021; Gan et al., 2019). Knowing that *Tnni3k* partially contributed to the end-state frequency of MNDCMs, we hypothesized that *Tnni3k* also plays a role in the ploidy dynamics observed after P21 in A/J mice. To test this, we followed *Tnni3k* global knockout mice (*Tnni3k^−/−^*) compared to wildtype littermates (*Tnni3k^+/+^*) maintained on a C57Bl/6J background for ploidy composition (Figure 3A) and total cardiomyocyte numbers (Figure 3B). At P21, MNDCM frequency was not different across genotypes, both fell below 2% (Figure 3A). From 3-6 weeks of age, *Tnni3k^+/+^* MNDCM numbers stayed constant around 1-1.5%. However, MNDCM frequency in *Tnni3k^−/−^* mice gradually increased over this same time course (P=0.02). At early adulthood, *Tnni3k* knockout mice had three times more MNDCMs than wildtype littermates (P=0.009), but this was still 3-fold less than typically observed in A/J mice, consistent with previous reports. Total cardiomyocyte number did not increase between P21 and 6-week timepoints in knockout animals, nor did it differ across genotypes (Figure 3B). When we repeated the experiment from Figure 2B (daily EdU injections from P14-P20) in these animals, there were fewer total EdU-positive cardiomyocytes in *Tnni3k^−/−^* mice (Figure 3C, P=0.0006), consistent with what we observed in A/Js by this same method (Figure 2D). Strikingly, at 6 weeks, we saw significantly more EdU-positive MNDCMs in *Tnni3k*^−/−^ compared to *Tnni3k^+/+^* mice (Figure 3D, P=0.0071). However, comparing *Tnni3k^−/−^* 6-week preparations to P21, we observed only a trending increase in EdU-positive MNDCMs (Figure 3D). Further, the frequency of EdU-positive MNDCMs were even more rare than observed in A/J with this same injection regimen (Figure 2E). Taken together, these data suggest that *Tnni3k* hypomorphism is only partially responsible for the ploidy phenotypes observed in A/J mice.

**Figure 3.**
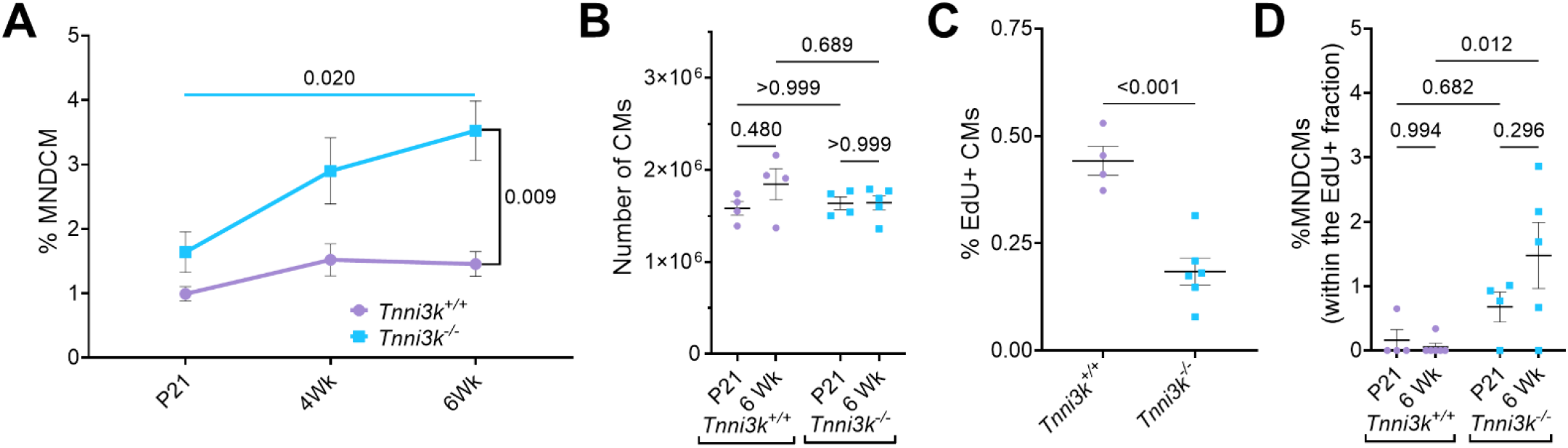
*Tnni3k* ablation in C57Bl/6J partially phenocopies A/J ploidy dynamics. **(A)** Percent MNDCM over time in *Tnni3k^−/−^* vs *Tnni3k^+/+^* (maintained on a C57Bl/6J background). **(B)** Number of cardiomyocytes counted via hemocytometer after Langendorff dissociation at P21 and 6 weeks (N=4-5). **(C)** Percent EdU+ cardiomyocytes at P21 following P14-P20 daily EdU injections. **(D)** Quantifications of EdU-positive MNDCMs represented as a percent of total EdU-positive cardiomyocytes in *Tnni3k^−/−^* and *Tnni3k^+/+^* at P21 and 6 weeks.

### A genetic locus, including Runx1, associates with MNDCM frequency

To identify additional genetic contributors to ploidy dynamics observed in A/J mice, we returned to the genome-wide association study (GWAS) which mapped frequency of mononuclear cardiomyocytes across the hybrid mouse diversity panel (HMDP) published by ((Patterson *et al*., 2017). With any genetic resource, including the HMDP, relevant genetic loci may be masked by complex gene-gene interactions including additive, suppressive, or epistatic relationships. Within the HMDP there are 4 panels of recombinant inbred (RI) strains: BXD (C57Bl/6J x DBA/2J), AXB/BXA (A/J x C57Bl/6J, and vice versa), BXH (C57Bl/6J X C3H/J), and CXB (Balb/cJ X C57Bl/6J); inclusion of any one of the RI panels could suppress or washout a relevant locus. With this logic, we re-ran the original phenotypic data, frequency of mononuclear cardiomyocytes across 120 strains, but excluded the BXD panel (44 strains), and a locus on chromosome (Chr) 16 rose in significance (Figure 4A versus Figure 4B (Patterson *et al*., 2017)). With the BXD panel removed, the AXB/BXA RI panel makes up 35% of the remaining data (27 AXB/BXA strains of 76 total remaining strains after BXDs have been removed); thus, it is a major contributor to the statistical significance. Interestingly, when the AXB/BXA panel is instead removed from the phenotype data, the locus is completely lost (Figure 4C). Taken together, these observations indicate that the Chr16 locus associated with the frequency of mononuclear cardiomyocytes was dependent on polymorphisms between A/J and C57Bl/6J.

**Figure 4.**
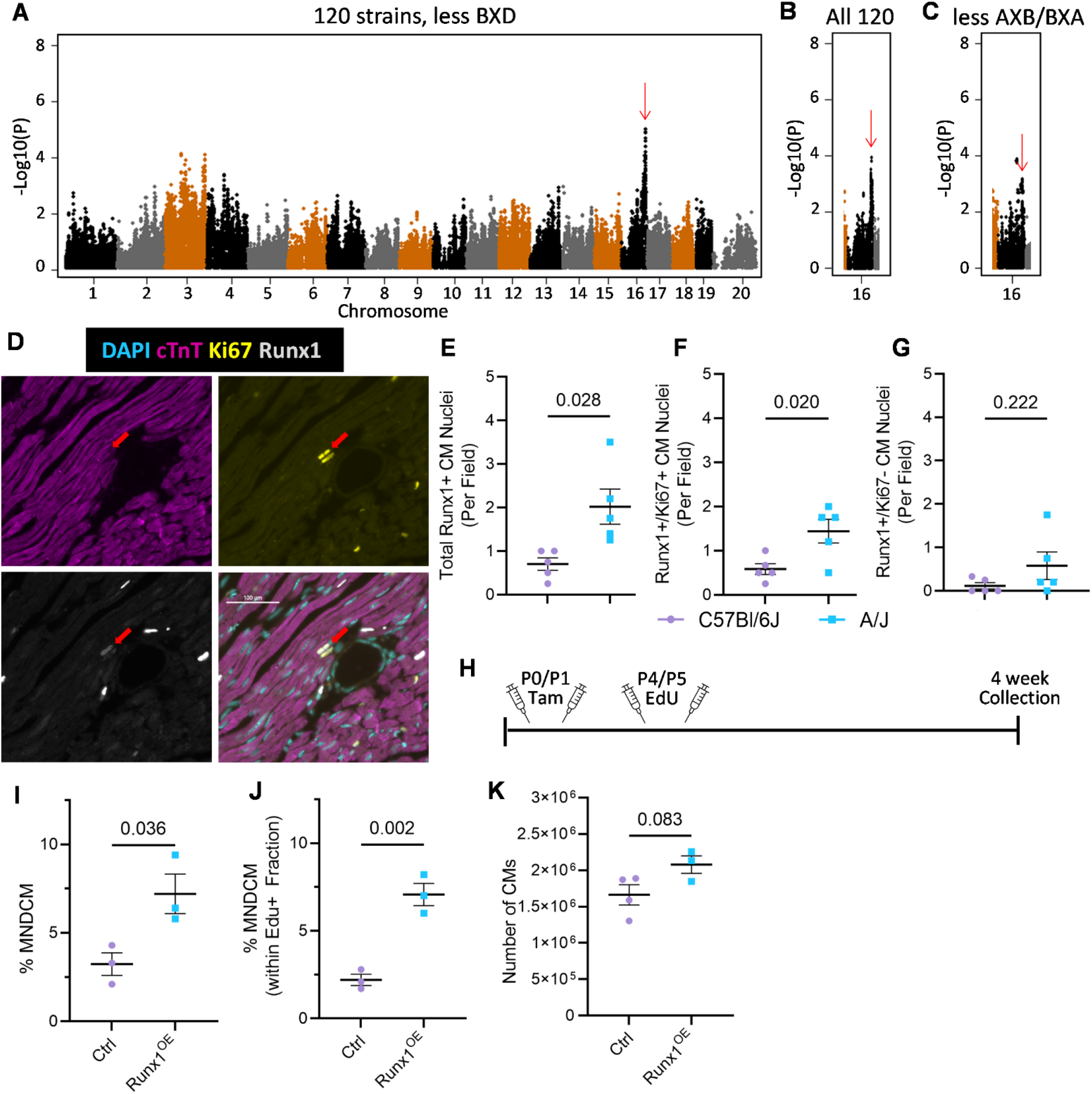
*Runx1* overexpression in C57Bl/6J is sufficient to induce A/J-like ploidy phenotypes. **(A)** Manhattan plot for genome association utilizing phenotypic data collected by (Patterson *et al*., 2017) after removing the BXD RI panel. **(B)** Manhattan plot for genomic association within Chr16 with all 120 strains included. **(C)** Manhattan plot for genomic association within Chr16 after removing the AXB/BXA RI panel. **(D)** Representative fluorescent image of a Runx1 (greyscale), Ki67 (yellow), cTnT (magenta), and DAPI (cyan). Red arrowhead points to a Runx1, Ki67-double positive cardiomyocyte. Scale bar = 100uM. **(E)** Quantification of percentage of Runx1-positive cardiomyocyte nuclei per 20x field. All pictures were taken in the left ventricle at P21 comparing C57Bl/6J to A/J **(F)** Quantification of Runx1, Ki67-double positive cardiomyocyte nuclei per 20x field. **(G)** Quantification of Runx1-positive, Ki67-negative cardiomyocyte nuclei per field. **(H)** Timeline depicting tamoxifen and EdU injections as well as collection timepoints for Runx1 OE Experiments 1 (Exp 1, top) & 2 (Exp 2, bottom). **(I)** Percent MNDCM at 4 weeks of age across *Runx1* OE and Cre-positive control (Ctrl) littermates following Exp 1 injection protocol. **(J)** Quantification of Edu-positive MNDCM as a percent of total EdU-positive cardiomyocytes following Exp 1 injection protocol. **(K)** Quantification of total cardiomyocyte number by hemocytometer following Exp 1 injection protocol.

Based on criteria set by (Wang et al., 2016) for determining locus range, the identified Chr16 locus spans approximately 4.2 mega base pairs (Mbps): 89,459,997 *Tiam* – 93,665,575 *Dop1b*. In another study using the HMDP, we used reduced representational bisulfite sequencing to identify changes to CpG methylation sites that affect cardiac phenotypes. Using the same BXD-removed data which identify the genomic locus, we performed an epigenome-wide association study (EWAS) and identified a locus-wide significant (P=3.9E-4) association between the methylation status of a CpG within the locus and the percent mononuclear cardiomyocytes. Leveraging the significantly smaller block sizes of EWAS loci (Orozco et al., 2015), we were able to narrow down the Chr16 locus further to approximately 92,763,000 – 93,665,500. Within this refined locus are only six protein coding genes: *Runx1, 1810053B23Rik, Setd4, Cbr1, Cbr3*, and *Dop1b*. Of these six genes, we identified *Runx1* as an interesting candidate as it has been used as a marker of cardiomyocyte regression to a less differentiated state prior to cell cycle re-entry (D’Uva et al., 2015; Kubin et al., 2011). In addition, Runx1 has been shown to interact with Yap (Chuang and Ito, 2021), a transcription factor well established for its regulation of cardiomyocyte proliferation (Monroe et al., 2019). With this in mind, we first asked if Runx1-positive cardiomyocytes were more prevalent in A/J versus C57Bl/6J hearts at P21, just preceding expansion of the MNDCM population in A/Js. Knowing that DNA synthesis was not different across strains at this timepoint, we instead combined Runx1 antibody stain with a more general cell cycle marker, Ki67 (Figure 4D). Interestingly, A/J ventricles not only displayed almost 3-fold more Runx1-positive cardiomyocytes than C57Bl/6J ventricles (Figure 4E), but Runx1-positive cardiomyocytes in A/J ventricles were more frequently double positive for Ki67 than C57Bl/6J cardiomyocytes (Figure 4F). Very few Runx1-positive Ki67-negative cardiomyocytes were found in either strain and there was no difference across genotypes in this population (Figure 4G).

### Runx1 overexpression is sufficient to induce cytokinesis and expansion of the MNDCM population in C57Bl/6J

To directly test Runx1’s ability to induce A/J-like phenotypes in C57Bl/6J mice, we utilized a conditional overexpression allele whereby *Runx1* cDNA preceded by a *LoxP-STOP-LoxP* cassette was inserted into the ubiquitous *Rosa26* locus [G(*tfRosa)26*^tm1(RUNX1)Mα^, referred to throughout as Runx1^OE^] (Qi et al., 2017; Yzaguirre et al., 2018). When crossed to the *Myh6-MerCreMer* driver (JAX Stock 011038), these mice overexpress *Runx1* only in cardiomyocytes following tamoxifen induction. Both alleles have been backcrossed and maintained on a C57Bl/6J background in our laboratory. Mice with a single copy of the *Runx1* transgene were crossed to mice with two copies of the *Cre* transgene, resulting in litters with either *Myh6-MerCreMer*^+/−^ only (Ctrl) or *Myh6-MerCreMer*^+/−^; *Rosa26*^tm1(RUNX1)Mα^ (Runx1^OE^). All animals received tamoxifen, which was administered with two subcutaneous injections at P0 and P1 (Figure 4H). With this injection regimen, we observed that most cardiomyocytes expressed Runx1 (Supp Figure 4A). Animals were also injected with EdU at P4 and P5 and hearts were collected at 4 weeks of age as a single cell suspension for nucleation and ploidy analysis (similar to Exp #1, Supp Figure 2A). First, considering all cardiomyocytes regardless of EdU status, Runx1^OE^ mice displayed twice as many MNDCMs compared to Ctrl control littermates (Figure 4I). There was no measurable change in the number of EdU-positive cardiomyocytes with this injection regimen (Supp Figure 4B), however we did observe a significant increase in the number of EdU-positive cardiomyocytes MNDCMs, suggesting they had completed cytokinesis (P=0.002, Figure 4J). Further, this resulted in a trending increase in total cardiomyocyte number, from 1.66M in Ctrl hearts to 2.08M in Runx1^OE^ hearts (P=0.08, Figure 4J). Taken together, these data indicate that *Runx1* is sufficient to induce A/J-like ploused reduced representational bidy phenotypes in C57Bl/6J mice.

### Single nucleus RNA sequencing characterizes distinct cardiomyocyte subpopulations for C57Bl/6J and A/J mice just prior to ploidy reversal

To identify and characterize the subpopulation of cardiomyocytes uniquely primed to expand within A/J hearts, we performed single nucleus RNA sequencing on 56,661 nuclei isolated from P21 hearts, just prior to the reversal event. Several past studies have enriched for cardiomyocyte nuclei by staining and sorting for Pcm1-positive nuclei prior to performing transcriptomics, however we have observed that as many as 13.0+/2.6% of adult A/J cardiomyocyte nuclei do not exhibit the classic perinuclear staining pattern and that there may be differential localization across ploidy classes (Supp Figure 5A-B). Therefore, we avoided using this enrichment strategy, and instead relied on post hoc gene expression to identify cardiomyocytes. We captured 28,070 total nuclei from 6 pooled C57Bl/6J hearts and 28,591 nuclei from 6 pooled A/J hearts. A median of 903 genes were mapped to each nucleus. Datasets were integrated from all samples using Seurat (Hao et al., 2021) and PHATE cell-type clustering (Moon et al., 2019) was subsequently run collectively, identifying 30 distinct clusters within the parent sample (Figure 5A). Individual cell types were identified by PHATE DimPlot ((Moon *et al*., 2019), Figure 5B) and further confirmed based on the top defining genes for each cluster. From this analysis clusters 7, 14, 21, and 24 were confidently identified as cardiomyocytes. These four clusters underwent doublet discrimination (McGinnis et al., 2019) and from this, 7,632 cardiomyocytes (4,240 C57Bl6/J and 3,392 A/J) were reclustered by Seurat (Hao *et al*., 2021), resulting in 12 cardiomyocyte clusters (Figure 5C). All 12 clusters express classic cardiomyocyte markers, including *Tnni3, Myh6*, and *Tnnt2*, (Supp Figure 5C).

**Figure 5.**
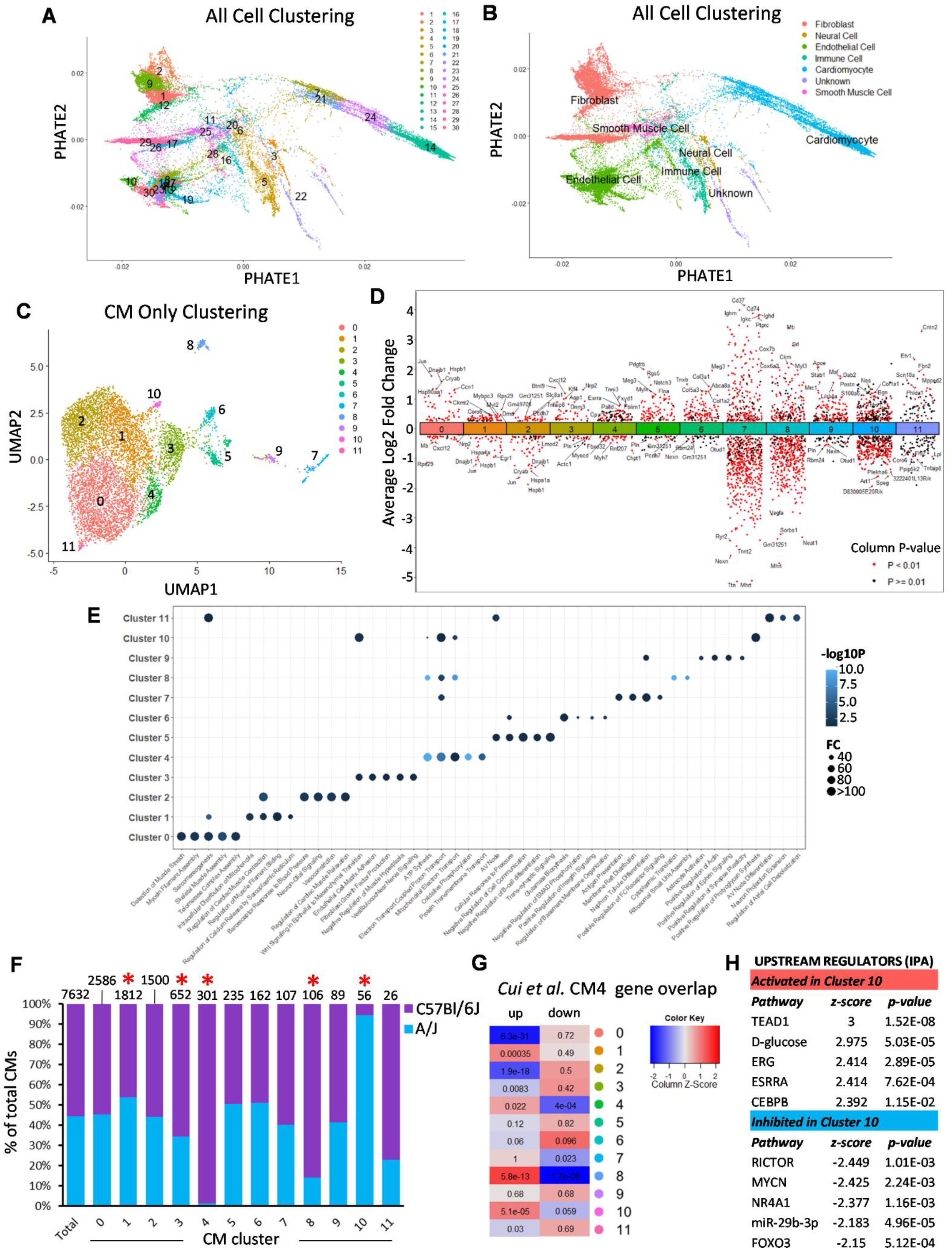
Single nucleus RNA sequencing identifies a unique cardiomyocyte subpopulation in A/J hearts. **(A)** Potential of Heat-diffusion for Affinity-based transition Embedding (PHATE) Dimplot of all 56,661 nuclei isolated from P21 A/J and C57Bl/6 hearts. **(B)** PHATE Dimplot identifying likely cell identifiers as determined by cellKb. **(C)** 7,632 nuclei from Clusters 7, 14, 21, and 24 from parent clustering plot reclustered by Uniformed Manifold Approximation Projection (UMAP) following doublet removal. **(D)** Dot plot of the uniquely upregulated and downregulated genes for each of the 12 cardiomyocyte clusters relative to other clusters. P<0.01 (red), P≥0.01 (black). **(E)** Top 5 Gene Ontology Terms represented by the upregulated gene lists for each cluster (P<0.01 and X Fold change). **(F)** Break down of number of A/J and C57Bl/6J nuclei represented in each cardiomyocyte cluster. Total number of nuclei for each cluster listed above bar. * indicates statistical deviation from the expected distribution (P<0.05 following Bonferroni correction) **(G)** Heatmap indicating the overlap of genes expressed by each cluster with the genes up and down regulated in Cui et al. (2020) CM4 cluster. Z-score normalized to column. **(H)** Top 5 activated and inhibited pathways identified by IPA upstream regulator analysis on the genes that define cluster 10.

To generally classify each cardiomyocyte cluster, we identified significantly enriched or depleted genes in each cluster (FDR-corrected P<0.01, 1.25-fold change, Figure 5D). The cluster-specific enriched genes (P<0.01, 1.5-fold change) were then also run through Panther Gene Ontology (GO) analysis (Figure 5E, (Thomas et al., 2022)). The majority of cardiomyocyte nuclei fell in clusters 0, 1, and 2, which were largely characterized by maturation processes like sarcomere organization, filament assembly and sliding, mitochondrial distribution, calcium ion signaling, and contraction-relaxation. Clusters 4 (301 nuclei) and 8 (106 nuclei) were predominantly derived from C57Bl/6J hearts, while Cluster 10 (56 nuclei) was identified as being almost exclusively A/J (Figure 5F, Supp Figure 5D).

We first attempted to determine if any cluster was predicted to be in the cell cycle. Tricycle analysis (Zheng et al., 2022) identified only one cluster, Cluster 9 (P=3.36E-04), however this cluster appears to be defined by many genes traditionally associated with leukocyte lineages (*Cyth4, Fyb, and Csf1r*, Figure 5D) and thus may represent a cluster of doublets that did not reach doublet cutoff criteria. Having not identified any clusters considered to be “cycling” by traditional means, we instead looked for gene expression overlap with cluster “CM4” from Cui et al. (2020) which identified CM4 as being a proliferative population unique to regenerative hearts (Cui et al., 2020). Notably, only two clusters from our data, Clusters 4 and 8, were identified as being significantly enriched for genes upregulated in Cui et al CM4 and significantly depleted for genes downregulated in Cui et al. CM4 (Figure 5F). Cluster 10, predominantly made up of A/J cardiomyocytes, was also significantly enriched for genes upregulated in Cui et al. CM4, but the depletion of downregulated genes was just above our FDR-corrected p-value range (P=0.059).

As we were interested in identifying a cluster unique to A/J hearts, we further investigated Cluster 10. We ran genes that were enriched in Cluster 10 relative to other cardiomocyte clusters (96 genes, P<0.01, 1.5-fold different from other clusters) through Ingenuity Pathways Analysis (IPA, *Qiagen*) and identified upstream regulators of gene networks. The top 5 predicted activated upstream regulators were *Tead1*, D-glucose, *Erg, Esrra*, and *Cebpb*, while *Rictor, Mycn, NR4A1*, miR-29b-3p (and other miRNAs with AGCACCA seed sequence), and *Foxo3* were predicted upstream regulators of inhibited gene networks (Figure 5G). The activated pathways are particularly interesting as *Tead1* is a transcription factor activated by Yap binding (Flinn et al., 2020). *Erg* is a ETS-family of transcription factor, and *Erg* and *Runx1* pathways are highly integrated in hematopoietic biology (Martens et al., 2012). In a recent study, both *Tead1* and *Erg (ETS*) binding motifs were identified by a Runx1 chromatin-immunoprecipitation (Gilmour et al., 2018), suggesting possible interactions between Runx1 and these transcription factors at their respective canonical DNA binding sites. Finally, *Cebpb* is considered a master regulator of cardiac hypertrophy (Bostrom et al., 2010), and has been linked to *Runx1* in both dorsal root ganglia (Ugarte et al., 2012) and early hematopoiesis (Lichtinger et al., 2012). Thus, at least 3 of the top five activated pathways are linked to *Runx1. Runx1* itself is also weakly identified by IPA upstream regulator analysis of cluster 10 defining genes (P=0.02) though directionality is not predicted.

## DISCUSSION

We initiated the current study with the goal of determining if mice with divergent cardiomyocyte ploidy distribution in adulthood arrive at these end states via distinct developmental pathways. In line with previously published literature (Alkass *et al*., 2015; Soonpaa *et al*., 2015), we confirmed that C57Bl/6J cardiomyocytes reach their terminal composition in a linear manner. By P14, cell cycle activity in C57Bl/6J cardiomyocytes is largely complete, and ploidy remains static thereafter. In contrast, A/J mice achieve their end state by a much more dynamic process. A/J cardiomyocyte polyploidy peaks around P21, after which there is a substantial expansion of the 1×2N population. Between P21 and 6-weeks of age, ploidy equilibrates, and A/J hearts arrive at their final adult frequency of MNDCMs which is more than 5x higher than observed in C57Bl/6J hearts. Concomitant with this expansion of MNDCMs in A/J, there is an increase in total cardiomyocyte numbers not observed in C57Bl/6J. Using single cell suspension methods paired with EdU labeling, we were able to identify delayed cytokinetic events in A/J, which again were not observed in C57Bl/6J. Further, there was a quantifiable decrease in polyploid cardiomyocytes, suggesting the expanding MNDCM pool is arising from a polyploid source. We believe this to be the first observation of ploidy reversal in cardiomyocytes, a phenomenon first reported in the liver field (Duncan *et al*., 2010). The divergent processes observed when comparing two inbred mouse strains suggest a knowledge gap remains regarding cardiomyocyte polyploidization and maturation, necessitating further experimentation on the unique phenotypic and genetic profiles across strains.

Through our examination of this MNDCM expansion unique to A/J hearts, we did not detect measurable amounts of DNA synthesis after P21, suggesting MNDCM expansion was not achieved through canonical mitotic means (i.e. a residual MNDCM population simply proliferated through the traditional mitotic cell cycle), but rather cell cycle completion from a pre-existing polyploid cell occurred. This led us to investigate the potential for delayed cytokinesis. Some have reported a synchronous proliferative burst of cardiomyocytes restricted to a narrow window of time during postnatal development (Naqvi et al., 2014). While we did not observe such an expansion of cardiomyocytes in C57Bl/6J, it remains possible that we missed an equivalent narrow window of cell cycle activity in A/J due to our experimental design.

The evaluation of cytokinesis is limited by insufficient available experimental strategies (Auchampach *et al*., 2022). Traditional markers, like Aurora Kinase B (AKB) notoriously localize to the midbody in both cardiomyocytes undergoing cytokinesis and those that ultimately fail to complete cytokinesis and become multinucleated instead (Engel et al., 2006; Leone et al., 2018). Thus, on its own AKB does not distinguish cytokinesis. Other strategies like the *Mosaic Analysis with Double Markers* (MADM) mouse (Ali et al., 2014) and sparse labeling followed by clonal analysis have been successfully used by several groups to confirm cell division (Bradley et al., 2021; Liu et al., 2021), but require engineered alleles to be bred into the experimental model. This was not practical in our case where we hoped to compare two divergent genetic backgrounds. We instead employed a universally applicable strategy, which combines EdU labeling with Langendorff single cell suspensions. With this method we were able to identify cells that had definitively completed cytokinesis in A/J hearts by 4 or 6 weeks, which were not present at P21. Further, this methodology allowed us to powerfully examine other ploidy classes and infer ploidy dynamics, which could not be explored by any of the histological methods described above. Admittedly, analysis of the ploidy classes by our method is largely based on ratios, and we have not excluded the potential of cell death within specific cardiomyocyte classes which would shift these ratios. However, a general increase in the number of total cardiomyocytes after P21 indicates that this possibility is unlikely.

Our analysis of genetically divergent inbred mouse strains in a controlled laboratory setting suggests a genetic component to the regulation of polyploidization. We sought multiple strategies to identify possible mechanisms. First, a previously performed GWAS on frequency of mononuclear cardiomyocytes (Patterson *et al*., 2017) provided a source of potential candidates that may reciprocally influence the unique developmental ploidy dynamics observed in A/J mice. *Tnni3k*, arising from this GWAS, has a naturally occurring polymorphism and has been previously demonstrated to control the size of the MNDCM population (Patterson *et al*., 2017). The variant is an alternative donor “G” four base pairs away from the consensus exon 19-20 splice site, resulting in a frame shift and premature stop codon (Wheeler *et al*., 2009). Ultimately, this mutation acts as a hypomorphic allele, and is carried by A/J mice, while C57Bl/6J carry the wild-type “A” at this locus. Here, we show that a genetically engineered loss-of-function mutation on the C57Bl/6J background partially phenocopies A/J cardiomyocyte ploidy progression. We see similar expansion of the MNDCM population between P21 and 6 weeks, as well as reduced DNA synthesis, and increased completion of cytokinesis post wean. However, the phenotypes in the engineered *Tnni3k* null C57Bl/6J mice are modest in comparison to A/J, suggesting polygenic contribution.

A second locus from the GWAS on Chr16 included *Runx1*, which has been implicated in the reversion of cardiomyocytes to a less differentiated state (D’Uva *et al*., 2015; Kubin *et al*., 2011; Zhang et al., 2016). Unlike *Tnni3k*, no obvious protein coding mutations distinguishing A/J from C57Bl/6J mice within this gene. We employed epigenome-wide association analysis to identify differential methylation status within the Chr16 locus and found that the *Runx1* transcriptional start site was differentially methylated between the two strains prompting us to select *Runx1* as a candidate gene in the locus. Indeed, we found more cardiomyocytes expressing *Runx1* in A/J compared to C57Bl/6J hearts at P21. Overexpression of *Runx1* in a C57Bl/6J background demonstrates that *Runx1* expression alone is sufficient to induce A/J-like cardiomyocyte ploidy phenotypes on the C57Bl/6J genetic background. These observations could prompt exploration of the role of *Runx1* in regeneration contexts. It is worth noting, however, that while we landed on *Runx1* from the Chr16 locus for a variety of reasons, there are 38 protein coding genes within the 4.2Mbps locus, and four other genes within the locus have protein coding variants between A/J and C57Bl/6J: *Ifnar1, Gart, Son*, and *Setd4*. Of the four genes, only the variant of *Setd4* is predicted to be “possibly damaging” by PolyPhen (Adzhubei et al., 2010) and SIFT (Ng and Henikoff, 2001) while the others are predicted to be benign. It remains possible that *Setd4* is a gene worth exploring in this context.

Finally, we performed single nucleus transcriptomic analysis in attempt to identify a unique subpopulation of cardiomyocytes in A/J hearts that might represent a population capable of ploidy reversal. While we cannot be certain that Cluster 10 represents the unique population responsible for the observed ploidy reversion, it did unveil molecular pathways which appear to be uniquely activated in a subset of A/J cardiomyocytes. *Tead*, the top hit to come from this analysis, is quite striking as an established effector of the Hippo-Yap pathway (Flinn *et al*., 2020) and cardiomyocyte proliferation (Monroe *et al*., 2019). *Tead*, along with 2 other identified pathways, *Erg* and *Cebpb*, interact with *Runx1* in other cell types (Gilmour *et al*., 2018; Lichtinger *et al*., 2012; Ugarte *et al*., 2012) lending support to the hypothesis that endogenous Runx1 expression in A/J hearts contributed to cardiomyocyte ploidy phenotypes. Further exploration of these pathways in conjunction with one another is warranted.

The function of polyploidization in cardiomyocytes remains unclear. Some have suggested developmental polyploidization, through the increase in total cellular DNA, supports enhanced rates of biosynthesis for production of contractile apparatus. Others hypothesize it promotes cell cycle exit as a strategy for both energy preservation and for preventing the disassembly of the pseudosyncitium inherent to cell division. In disease contexts, many in the field postulate that MNDCMs are uniquely capable of mounting a regenerative response based on the observation that polyploidization coincides with loss of regenerative competence. This hypothesis is supported by several recent studies where ploidy phenotypes were manipulated or altered and regeneration in turn was impacted (Gonzalez-Rosa *et al*., 2018; Han *et al*., 2020; Hirose *et al*., 2019; Patterson *et al*., 2017). Additionally, high levels of polyploidy have been observed in various cardiomyopathies and heart failure (Beltrami et al., 1997; Brodsky et al., 1994; Gilsbach et al., 2018). Despite these numerous studies, further work is warranted to determine what effect either blocking or driving the polyploidization process has on heart function. Understanding the genetic mechanisms that regulate polyploidization may help us better understand its role in normal cardiac physiology, myocardial regeneration, and heart failure.

## ACKNOWLEDGEMENTS

We thank Dr. Miao Cui for sharing her gene list from CM4. This work was supported by the American Heart Association, 18CDA34110240 awarded to M.P., and the National Institutes of Health, F31HL162468 awarded to S.K.S, R01HL155085 awarded to M.P., F32HL150958 awarded to M.A.F, R01HL141159 awarded to C.C.O, and R00HL138301 awarded to C.D.R.

## AUTHOR CONTRIBUTIONS

Conceptualization, S.K.S., A.L.P., C.C.O, C.D.R., M.P.; Methodology, S.K.S., M.A.F., C.D.R., M.P.; Investigation, S.K.S., A.L.P., M.E.K., M.A.F., C.L., T.B., K.A.A., P.F., C.D.R., M.P.; Writing – Original Draft, S.K.S., C.D.R., M.P.; Writing – Review & Editing, S.K.S., A.L.P., M.A.F., T.B., K.A.A., P.F., C.C.O., C.D.R., M.P.; Funding Acquisition, S.K.S., M.A.F., C.C.O., C.D.R., M.P.; Supervision, C.C.O., C.D.R., M.P.

## STAR*METHODS

### Mice

All animal experiments were approved by and performed in accordance with the Institutional Animal Care and Use Committee of the Medical College of Wisconsin. A/J (JAX stock #000646) and C57Bl/6J (JAX stock #000664) mice were either purchased from Jackson Laboratory, Bar Harbor, Maine and allowed to acclimate for at least one week prior to experimental start or were bred inhouse from breeders originally purchased from Jackson Laboratory. *Tnni3k* global knockout mice were generated as described in (Patterson *et al*., 2017), G*t(Rosa)26*^tm1(RUNX1)Mα^ mice (Qi *et al*., 2017; Yzaguirre *et al*., 2018) were obtained from the Speck Laboratory at University of Pennsylvania, and *Myh6-MerCreMer* (JAX Stock 011038) mice were obtained from JAX Laboratory. All three alleles have been backcrossed and maintained at least 8 generations on a C57Bl/6J background by our laboratory. For all timed experiments, birth (postnatal day 0, P0) was assumed to have taken place at 12:00 AM. Only litters with 3-8 pups were used; and runts were excluded from any analysis. All experiments include a mix of virgin males and females, although sex was not determined for neonates <P21. In experiments where a mix of the sexes were used, no phenotypic differences between the sexes were observed. Animals were housed as compatible pairs or groups in ventilated cages on 12-hour light/dark cycles with *ad libitum* access to water and food. Euthanasia was performed in accordance with the recommendations of the American Veterinary Medical Association. Neonates ≤P14 were decapitated with surgical scissors, while animals ≥P21 underwent cervical dislocation following isoflurane-induced anesthesia.

### EdU Administration

5-ethynyl-2’-deoxyuridine (EdU, Thermo Fisher, E10187) was resuspended in DMSO at 100mg/mL to create a stock solution, which was aliquoted and stored at −20°C for no longer than 3 months. A fresh, never thawed aliquot of stock solution was further diluted to 1mg/mL with sterile PBS no more than 30-minutes prior to administration. Mice were injected once per day by intraperitoneal (i.p.) injection at 10mg/kg. Animals were euthanized at time indicated. For a complete list of administration methods broken down by experiment, please see Table 1.

**Table 1.**
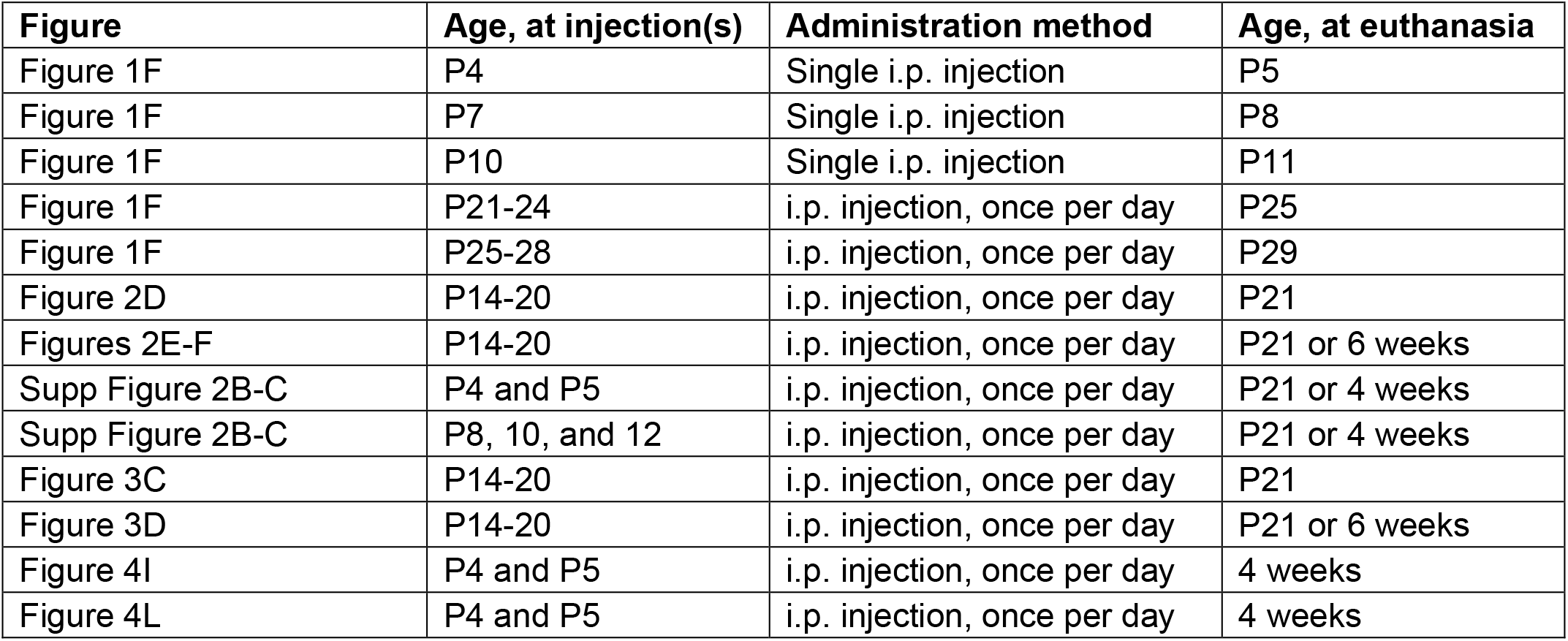
List of EdU injection strategies by experiment.

### Single cell ventricular suspensions

Hearts were extracted from euthanized mice by cutting the aorta just below the arch arteries, along with the other major vessels. Isolated hearts were washed in ice cold Kruftbruhe (KB) solution and secured by their aortas to a cannula of varying sizes (see Table 2) then tied off with a 3-0 silk suture. Atria were removed with Vannas micro spring scissors. Cannulated ventricles were then hung from a Langendorff apparatus and perfused with calcium-free Tyrodes buffer, followed by 1 mg/mL collagenase type II (Thermo Fisher, 17101015) dissolved in calcium-free Tyrodes buffer. Both solutions were warmed to 37°C. Volume of collagenase solution, along with size of cannula, varied by age of mouse (see Table 2 for details). Following perfusion, ventricular tissue was diced with dissection scissors, triturated in ice cold Kruftbrühe (KB) solution using a wide bore 1mL pipette, filtered through a 250μm mesh, and fixed by adding equal volume of 8% ice cold PFA and letting stand at room temperature (RT) for 10 minutes (final concentration of PFA = 4%). Filtering through the 250μM mesh was not used when assessing cardiomyocyte number. Following fixation, cell suspensions were spun down at 300G for 2 minutes and resuspended in PBS.

**Table 2.** Cannula size by mouse age.

| **Table 2. Cannula size by mouse age** |  |  |
| --- | --- | --- |
| Age of mouse | Cannula size | Volume of collagenase (1mg/ml) |
| P1 | 27 gauge | 15-20 mL |
| P4-9 | 27 or 22 gauge | 20-25 mL |
| P10-20 | 22 or 20 gauge | 25 mL |
| P21 and older | 20 or 18 gauge | 25-50 mL |

### Quantification of total cardiomyocyte numbers

Fixed unfiltered cells resultant from a whole heart (less atria) Langendorff digestion, were resuspended in 2 mL of PBS. While fully resuspended, a 20μl aliquot was drawn up with a 100 μl pipette and diluted 1:5 in PBS. This was repeated three separate times for each heart. Each of the three aliquots was counted in triplicate on a hemocytometer. Cardiomyocytes were distinguished from non-cardiomyocytes by size and morphology. All counts (9 in total) were averaged together to come up with the best possible estimate of total cardiomyocyte numbers. A two-way ANOVA with multiple comparisons (age and strain being the two dependent variables) and Tukey posthoc test were run to calculate the statistical significance of any inter- and intra-strain differences.

### Immunostain for single cell suspensions

Fixed ventricular cell suspensions were blocked with 10% normal goat serum (NGS, Thermo Fisher, catalog # 50062Z) and .01% triton X-100 for 1 hour at RT. Cells were incubated with primary antibody for either mouse anti-cTnT (1:500, Abcam ab8295), mouse anti-Actn2 (1:500, Sigma A7811), or rabbit anti-Pcm1 (1:400. Sigma Aldrich HPA023370) in blocking solution overnight at 4 °C. Cells were then washed twice in PBS with spins at 300G for 3 mins in between and incubated with Alexa Fluor 488 goat anti-mouse secondary (1:500, Thermo Fisher, A11029) in PBS for one hour at RT. During the last ten minutes of secondary incubation 4’,6-diamidino-2-phenylindole (DAPI 1mg/mL, 1:1000) was added to the suspensions. Cells were washed two times in diH_2_O and spun one final time. Cells from the final pellet were resuspended with Prolong Gold (Thermo Fisher, P26930), pipetted across a slide and cover slipped.

In the case of “cytokinesis” experiments as described by Figure 2A, cell suspensions were also stained with a Click-it EdU kit Alexa Fluor 555 (Thermo Fisher, C10339) according to the manufacturer’s protocol. This was performed after blocking and primary antibody but prior to addition of secondary antibody. The entire pellet was pipetted across slides at a density of ~15μL of pelleted cells per slide mixed with 20μL of Prolong Gold (Thermo Fisher P26930) and cover slipped.

### Ploidy analysis

Following staining, cardiomyocyte nucleation was quantified on a Nikon Eclipse 80i fluorescent microscope with a 20x objective. 300 healthy cardiomyocytes were counted for their nucleation (i.e. mono-bitri- or tetranucelated); cardiomyocytes with a spherical shape or frayed edges (accounting for less than 5% of any preparation) were excluded for being dead or dying. Additionally, fluorescent images were taken at 10x magnification with a Panda PCO camera and analyzed on NIS Elements software. For each animal, the nuclei from ~500 cells were evaluated for nuclear ploidy by calculating the sum DAPI intensity of each nucleus and normalizing it to a known 2N population, typically Tri and Tetranucleated cardiomyocytes. Nuclear ploidy was calculated separately for each nucleation class and the two independent measurements were combined to estimate the frequency of each ploidy class (i.e. 1×2N, 1×4N, 1×8N, 1×16N, 2×2N, 2×4N, 2×8N, Tri – 2×2N + 1×4N or 2×4N + 1×8N, and Tetra – 4×2N or 4×4N) represented as a percent of total. Because all ploidy subpopulations add up to 100% and are therefore interdependent on one another, a multivariate ANOVA with LSD posthoc test was used to compare inter- and intra-strain differences.

An estimate of total number of cardiomyocytes for each ploidy class, as in Figure 2G was calculated by multiplying the ploidy class percentages of each individual by the average of total cardiomyocyte number at each specific timepoint. Changes in cardiomyocyte numbers, as in Figure 2H, were calculated by subtracting the estimated total of a given ploidy class at 3 weeks from the estimated total of the same ploidy class at 4 weeks. Statistical significance in Figure 2H was assessed by multivariate ANOVA comparing 3- and 4-week timepoints.

### Cytokinesis analysis by single cell suspension

As with ploidy analysis, slides were scanned in their entirety on a Nikon Eclipse 80i fluorescent microscope with a 20x objective until at least 175 EdU-positive cardiomyocytes were counted for their nucleation. Total percentages of EdU-positive cardiomyocytes in the heart were estimated using the number of EdU-positive cardiomyocytes found on a single slide multiplied by the number of slides generated following staining and divided by the total number of cardiomyocytes quantified by hemocytometer for that same animal. While scanning, images of EdU-positive cardiomyocytes pictures were taken at 10x. EdU-positive cardiomyocyte nuclei were analyzed in NIS elements for DAPI intensity and normalized to a known 2N nucleus as above. At least 25 mononucleated cardiomyocytes were analyzed for each animal. A two-way ANOVA with multiple comparisons was run to show inter and intra strain differences.

### Histology

As above, hearts were dissected from euthanized mice, washed in KB solution until no longer beating, and hung on a Langendorff apparatus. Hung hearts were first perfused retroaorticaly with 5 mL of calcium-free Tyrodes to flush out any remaining blood, followed by 5 mL of ice cold 4% PFA and then further fixed in 4% PFA overnight at 4 °C (~12-18 hours). After washing three times in PBS, hearts were stored in 70% ethanol until further processing could take place. Briefly, hearts were dehydrated by progressive introduction of ethanol (80%, 90%, 100%) and cleared with xylene prior to being embedded in paraffin wax. Embedded tissues were sectioned from apex to outflow track (2-chamber view) on a Thermo Microm HM 355S microtome at 4μm thickness. Tissues were collected every 200-400 μm depending on the age of the animal/size of the heart.

### In situ fluorescence

Tissue sections were rehydrated by sequential introduction to ethanol solutions for 2 minutes each (Xylene, 100%, 90%, 80%, 70%, H2O) and heated at 100°C in Sodium Citrate buffer with 0.1% Triton-X-100 and .05% Tween-20, pH 6.0 for 30 minutes for antigen retrieval. After washing in PBS, slides were blocked with 5% normal donkey serum (NDS, Jackson ImmunoResearch 017-000-121) and 5% bovine serum albumin (BSA, VWR 97061-420) in PBS for 1 hour at RT. Primary antibodies, goat anti-NKX2.5 (1:250, Abcam ab106923), mouse anti-cTnT (1:500, Abcam ab8295), rabbit anti-Runx (1:250, Abcam ab209838), and/or rat anti-Ki67 (1:250, Invitrogen 14-5698-82) were diluted in blocking buffer and incubated on tissue sections at 37 °C for two hours in a humid chamber. Slides were washed in PBS and incubated with secondary antibodies Alexa Fluor donkey anti-goat 555 (Thermo Fisher A32816), donkey anti-mouse 647 (Thermo Fisher, A31571), donkey anti-rabbit 555 (Thermo Fisher A31572), or donkey anti-rat 488 (Abcam ab150153) respectively, diluted in PBS for one hour RT. Slides were washed in PBS and EdU incorporation was labeled using Click-it EdU Alexa Fluor 488 kit (Thermo Fisher, C10339) according to manufacturer’s protocol. After washing in PBS, tissues were incubated with 0.03% Sudan black B (SBB) dissolved in 70% ethanol for 20 minutes at RT followed by a PBS wash. DAPI was labeled using 10ug/ml in PBS for 5 mins at RT, followed by PBS wash. Slides were cover slipped using Prolong Gold (Thermo Fisher, P26930) and allowed to dry in the dark. Pictures were taken with a PCO Panda camera using a 20x objective on a Nikon Eclipse 80i fluorescence microscope. ~6 pictures were taken per animal in randomly selected regions throughout the left ventricle and the septum (equal representation of each). EdU-positive Nkx2.5-positive cardiomyocyte nuclei were quantified as a percent of total Nkx2.5-positive cardiomyocyte nuclei using NIS Elements software. Runx1 and Ki67 positive cardiomyocyte nuclei were identified by intersection with cTnT and represented as positive nuclei per region. A student’s T-test was run between strains at each time point or between strains at a single time point.

### Nuclear isolation for single nucleus RNA sequencing

On two occasions for each strain, hearts were excised from 3 littermates collected at P21. If 2 females and 1 male were used for the first collection, then the reverse was done on the second collection, such that the final sequencing represents 6 hearts, 3 males and 3 females. Excised hearts, with atria removed, were Langendorff perfused with 25 mLs of 1mg/mL collagenase as described above. Digested hearts were resuspended in ice cold KB and allowed to settle for 10 minutes on ice. We had hoped this would enrich for cardiomyocytes which are larger and heavier than other cell types, though it did not seem to help. Following 10 minute incubation, the supernatant was removed and the loose pellet was resuspended in 5mL of Lysis buffer prepared as described in (Cui and Olson, 2020) with only one adjustment – 50ul of 10% Triton-X-100 was added (final concentration 0.1%). Cells were incubated in Lysis buffer + Triton for 5 minutes on ice, after which they were homogenized with a Tissue Tearor electric tissue homogenizor (Model # 985370) at the second lowest setting for 20-30 seconds, and left to sit again for another 5 minutes on ice. They were then transferred through a 15mL glass dounce homogenizer and further homogenized with 20 strokes of the A pestle and 20 strokes of the B pestle. Homogenized cell suspensions were sequentially filtered through a 70uM, 40um, and 20um cell strainer to removed debris and undigested materials. Samples were then spun at 1000 G for 5 minutes and resuspended in 1mL of 2% BSA dissolved in d-PBS with RNaseOut (Invitrogen, 200U/mL). A small aliquot was set aside to serve as an unstained control for fluorescent activated cell sorting (FACS). The remainder of the suspension was stained with DAPI at 10ug/ml for 5 minutes on ice. Samples were spun at 1000G for 5 minutes and resuspended in fresh 2% BSA-RNaseOut solution.

Following staining, nuclei were sorted on a BD FACSMelody at 4°C. Following standard protocols, forward and side scatters were used to remove doublets. Unstained controls were used to set the V450 gate. 432,000 nuclei were collected into a 2mL centrifuge tube preloaded with 500uL of 2% BSA-RNaseOut solution. Sorted nuclei were spun down at 1000G for 5 minutes, supernatant was removed, and samples were resuspended in 100uL of 2% BSA-RNaseOut solution before proceeding to 10x library preparation.

### 10xGenomics cDNA and library preparation of nuclear samples

Sorted nuclei resuspended in a solution of D-PBS with 2% BSA solution and RNaseout (Invitrogen).

Nuclei were quantified with a Luna Fl cell counter (Logos Biosystems) and the volume was adjusted to obtain the ideal concentration of nuclei recommended by 10x Genomics (1000 nuclei/μL). Individual nuclei were paired with Chromium v3.1 gel beads and cDNA synthesis, barcoding, and dual index library preparation was performed using Chromium Next GEM V3.1 chemistry according to the manufacturer’s recommendation (10x Genomics). 10,000 nuclei were targeted for each sample with 13 cycles for cDNA amplification and13 cycles for sample index PCR. The fragment size of cDNA and libraries was assessed using Agilent’s 5200 Fragment Analyzer System to verify product quality prior to sequencing.

### Sequencing

4 libraries were sequenced at the Roy J. Carver Biotechnology Center at the University of Illinois, Urbana Champaign on a NovaSeq 6000 using one S4 lane with 2×150nt reads. Samples were demultiplexed and mapped to the mm10 genome using Cell Ranger v6.1.1 (10X Genomics). These data are accessible at BioProject using the ID PRJNA880279 in the Sequence Read Archive.

### Single nuclear RNA sequencing analysis

Each library was preprocessed using the Seurat 4.1.0 R package (Hao *et al*., 2021) to retain nuclei with unique feature counts between 200 and 2,500 and with fewer than 2% mitochondrial counts. The four libraries were then integrated into a single Seurat object using the Seurat functions FindIntegrationAnchors and IntergrateData, as previously described (Hao *et al*., 2021). Briefly, the functions act to perform batch correction on the data by identifying transformation vectors based on integration ‘anchors’ that show strong local ‘neighborhood’ correlation with similar cells, but a broad diversity of expression across the entire dataset. The functions leverage that information to merge cell clusters between samples together into an integrated whole. Doublets were identified and removed using DoubletFinder 2.0 (McGinnis *et al*., 2019), which generates artificial doublets from Seurat-processed clusters, inserts them into the original dataset and then identifies likely true doublets by identified cells with high correlation to numerous artificial doublets. Cell clustering was then performed using PHATEr (1.0.7) (Moon *et al*., 2019) and differentially expressed features identified using the FindAllMarkers function in Seurat. Differentially expressed genes were examined using cellKb 2.2.1 (*Biorxiv:* https://doi.org/10.1101/2020.12.01.389890) which compares the marker genes with nearly 40k marker gene sets representing 2,742 unique cell types across 10 species, by *Tabula Muris* (Tabula Muris et al., 2018), and cardiomyocyte-enriched clusters were separated for further analysis.

Cardiomyocyte clusters were re-clustered using the standard Seurat PCA-based pipeline and mapped with the UMAP projection. Differentially expressed genes were identified using the FindAllMarkers function as above, and gene ontology enrichments performed using Panther version 17.0 (Thomas *et al*., 2022) for both differentially expressed (FDR-adjusted P < 0.01, absolute value of fold change > 1.25) and highly expressed (FDR-adjusted P < 0.01, fold change > 1.5) genes in each cluster. Cell Cycle identity for each cell was determined using the tricycle transfer learning algorithm (1.2.1, (Zheng *et al*., 2022)) as described in the manuscript and enrichment of clusters for cells within the cell cycle determined by chi-squared testing.

Finally, the identifying genes from (Cui *et al*., 2020) CM4 were split into up- and down-regulated gene sets (absolute value of log2 fold change > 0.5 and FDR-corrected p-value < 0.001) and enrichment or depletion across the cardiomyocyte clusters was determined by Wilcoxon Man Whitney Correlation Corrected GSEA as implemented in the singleseggset R package (0.1.2.9, https://arc85.github.io/singleseqgset/index.html).

### Genome-wide association study

Phenotypes from 120 inbred mouse strains for the HMDP were taken from (Patterson *et al*., 2017). Averages for each strain underwent arcsin transformation to normalize the distribution of the data. Association testing was conducted on either 120 strains less the 44 strains of the BXD panel or on the 120 strains less the 27 strains of the AXB/BXA panel. Association testing of each single nucleotide polymorphism was performed in R software package as described in (Rau et al., 2015).

### Epigenome-wide association study

The data for this portion of the study comes from the control mice of a prior HMDP studying heart failure (Lahue, 2022; Rau *et al*., 2015). Briefly, DNA was isolated from left ventricles of 92 strains and sequenced using reduced representational bisulfite sequencing. DNA methylation was called using BSSeeker2 (Guo et al., 2013) using the mm10 genome build. Hypervariable CpGs were identified as CpGs which showed greater than 25% methylation variability in at least 10% of the studied strains as previously described (PMC4454894). Phenotypes were taken from (Patterson *et al*., 2017) as described above. EWAS was performed using the MACAU algorithm (Lea et al., 2015). Locus-wide significance was determined using the benjamini-hochberg correction. Locus boundaries were determined as previously described (Orozco *et al*., 2015).

## SUPPLEMENTAL INFORMATION

Please find Supplemental Tables 1 and 2, and Supplemental Figures 1-5 attached. Sequencing data have been uploaded to BioProject database (BioProject ID PRJNA880279).

