## Supplement for "Cardiomyocyte ploidy is dynamic during postnatal development and varies across genetic backgrounds"

### Cardiomyocyte ploidy dynamics during postnatal development vary between inbred mouse strains

This document includes:

Supplemental Table 1

Supplemental Table 2

Supplement to Figure 1

Supplement to Figure 2

Supplement to Figure 4

Supplement to Figure 5

**Supp Table 1.** Summary of descriptive statistics including N, Mean, and standard error of the mean (SEM), one-way ANOVA, and Tukey HSD post hoc analyses for Figure 1A.

| C57Bl/6J | | | |
|---|---|---|---|
| Age | N | Mean | SEM |
| P1 | 8 | 647,125 | 114,823 |
| P7 | 11 | 1,377,273 | 59,698 |
| P21 | 7 | 1,644,286 | 84,427 |
| 6Wk | 6 | 1,691,667 | 90,973 |

| A/J | | | |
|---|---|---|---|
| Age | N | Mean | SEM |
| P1 | 8 | 643,500 | 61,030 |
| P7 | 7 | 1,084,000 | 34,404 |
| P21 | 16 | 1,224,813 | 83,246 |
| 4Wk | 15 | 1,440,667 | 79,327 |
| 6Wk | 11 | 1,572,727 | 105,745 |

| **C57Bl/6J** | **ANOVA $p < 0.0001$** | |
|---|---|---|
| Tukey post hoc test | | |
| Timepoint | Timepoint | P-value |
| P1 | P7 | <0.0001 |
| | P21 | <0.0001 |
| | 6Wk | <0.0001 |
| P7 | P21 | 0.1331 |
| | 6Wk | 0.0768 |
| P21 | 6Wk | 0.9852 |

| **A/J** | **ANOVA $p < 0.0001$** | |
|---|---|---|
| Tukey post hoc test | | |
| Timepoint | Timepoint | P-value |
| P1 | P7 | 0.0414 |
| | P21 | 0.0003 |
| | 4Wk | <0.0001 |
| | 6Wk | <0.0001 |
| P7 | P21 | 0.8261 |
| | 4Wk | 0.0748 |
| | 6Wk | 0.0096 |
| P21 | 4Wk | 0.2584 |
| | 6Wk | 0.0300 |
| 4Wk | 6Wk | 0.7877 |

**Supp Table 2.** Multivariate ANOVA with Tukey HSD post hoc tests for Figure 1C.

| C57Bl/6J | Pillai's Trace P=0.00445 | | |
|---|---|---|---|
| Ploidy class | Timepoint | Timepoint | Tukey P-value |
| 2N | P7 | P14 | 2.24E-06 |
| | | P21 | 8.46E-07 |
| | | 6wk | 5.38E-07 |
| | P14 | P21 | 0.686 |
| | | 6wk | 0.382 |
| | P21 | 6wk | 0.942 |
| 4N | P7 | P14 | 0.001 |
| | | P21 | 5.87E-05 |
| | | 6wk | 6.82E-06 |
| | P14 | P21 | 0.263 |
| | | 6wk | 0.013 |
| | P21 | 6wk | 0.269 |
| 8N | P7 | P14 | 4.28E-06 |
| | | P21 | 3.80E-07 |
| | | 6wk | 8.69E-08 |
| | P14 | P21 | 0.073 |
| | | 6wk | 0.003 |
| | P21 | 6wk | 0.230 |
| 16N | P7 | P14 | 0.731 |
| | | P21 | 0.999 |
| | | 6wk | 0.408 |
| | P14 | P21 | 0.854 |
| | | 6wk | 0.951 |
| | P21 | 6wk | 0.570 |

| A/J | Pillai's Trace P=0.00337 | | |
|---|---|---|---|
| Ploidy class | Timepoint | Timepoint | Tukey P-value |
| 2N | P7 | P14 | 1.33E-07 |
| | | P21 | 1.01E-09 |
| | | 4wk | 1.26E-06 |
| | | 6wk | 6.36E-07 |
| | P14 | P21 | 0.592 |
| | | 4wk | 0.362 |
| | | 6wk | 0.380 |
| | P21 | 4wk | 0.007 |
| | | 6wk | 0.006 |
| | 4wk | 6wk | 1.000 |
| 4N | P7 | P14 | 0.989 |
| | | P21 | 0.737 |
| | | 4wk | 0.736 |
| | | 6wk | 0.999 |
| | P14 | P21 | 0.958 |
| | | 4wk | 0.954 |
| | | 6wk | 0.998 |
| | P21 | 4wk | 1.000 |
| | | 6wk | 0.783 |
| | 4wk | 6wk | 0.783 |
| 8N | P7 | P14 | 0.012 |
| | | P21 | 0.004 |
| | | 4wk | 0.160 |
| | | 6wk | 0.023 |
| | P14 | P21 | 1.000 |
| | | 4wk | 0.531 |
| | | 6wk | 0.944 |
| | P21 | 4wk | 0.379 |
| | | 6wk | 0.900 |
| | 4wk | 6wk | 0.859 |
| 16N | P7 | P14 | 0.971 |
| | | P21 | 0.619 |
| | | 4wk | 0.720 |
| | | 6wk | 0.076 |
| | P14 | P21 | 0.948 |
| | | 4wk | 0.977 |
| | | 6wk | 0.273 |
| | P21 | 4wk | 1.000 |
| | | 6wk | 0.538 |
| | 4wk | 6wk | 0.485 |

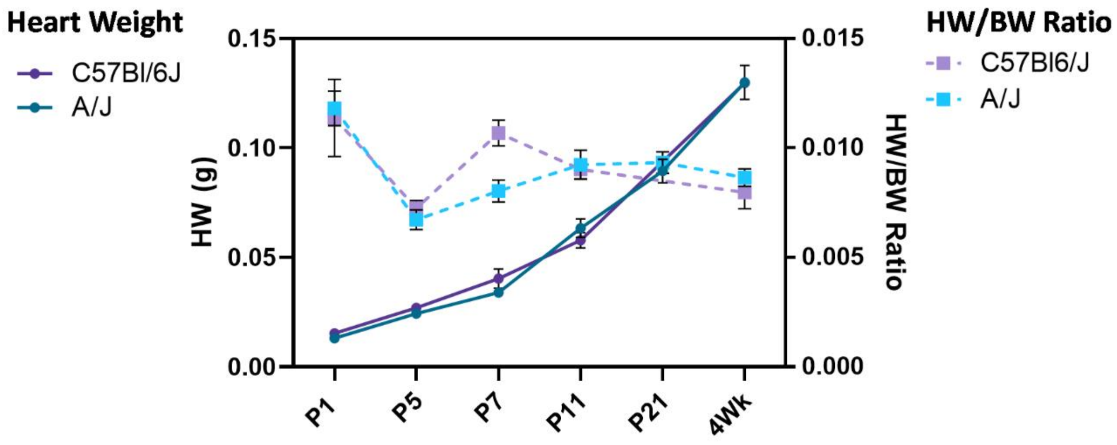


**Supplement to Figure 1.** Heart weight (HW, left y-axis) in grams (g) across A/J (light blue) and C57Bl/6J (light purple) at identified timepoints. HW to body weight (HW/BW, right y-axis) ratio across A/J (dark blue) and C57Bl/6J (dark purple) at identified timepoints. N = 3-17 animals. Error bars represent standard error of the mean.

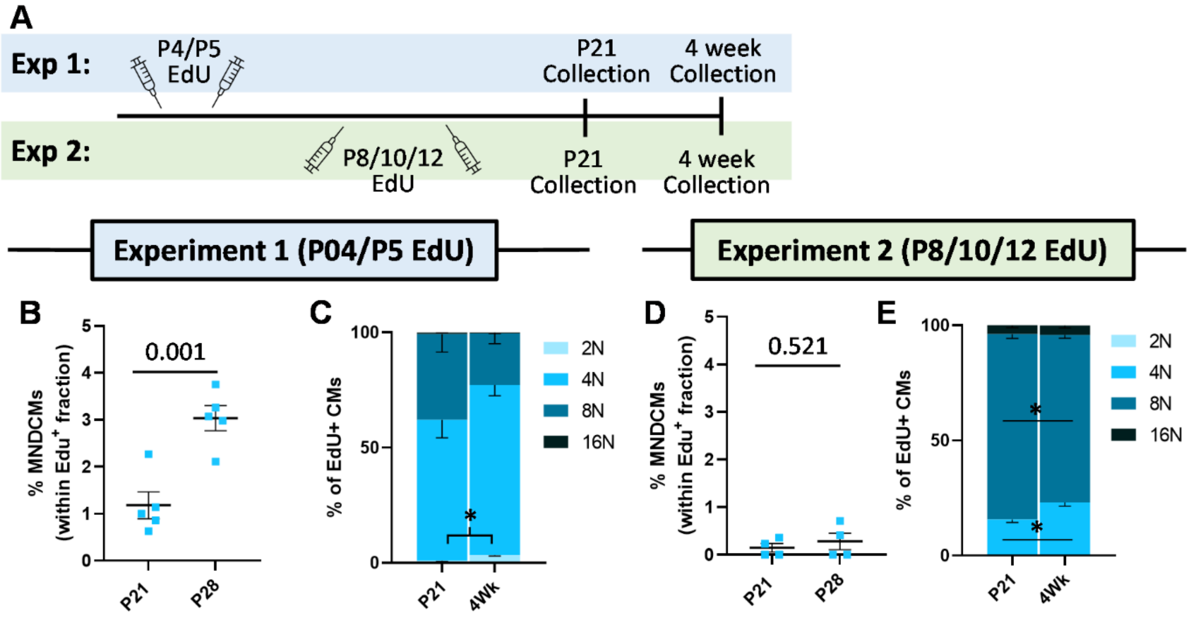


**Supplement to Figure 2. (A)** Schematic of two different EdU injection regimens on A/J mice for single cell suspension analysis of ploidy and cytokinesis. Experiment 1 (Exp 1, top) had single EdU injections on P4 and P5. Experiment 2 (Exp 2, bottom) had single EdU injections on P8, P10, and P12. **(B)** Quantifications of EdU-positive MNDCMs as a percent of total EdU-postive cardiomyocytes in A/J mice following Experiment 1 paradigm. **(C)** Quantifications of EdU+ cardiomyocytes broken down into total DNA content (i.e. 2N, 4N, 8N, or 16N) following Experiment 1 paradigm. * indicates P<0.05 (2N population). **(D)** Quantifications of EdU-positive MNDCMs as a percent of total EdU-postive cardiomyocytes in A/J mice following Experiment 2 paradigm. **(E)** Quantifications of EdU+ cardiomyocytes broken down into total DNA content (i.e. 2N, 4N, 8N, or 16N) following Experiment 1 paradigm. * indicates P<0.05 (4N and 8N populations).

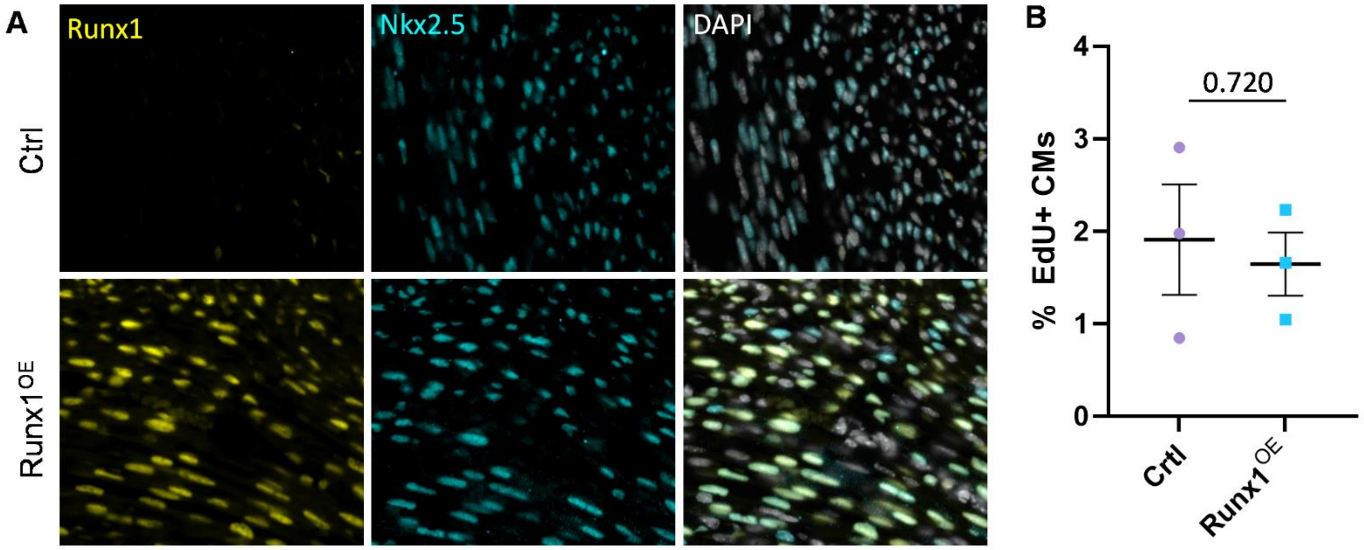


**Supplement to Figure 4. (A)** Immunofluorescent images for Runx1 (yellow), Nkx2.5 (cyan), and DAPI (greyscale) in Cre-positive Control animals (Ctrl) and Runx1$^{OE}$ hearts following two tamoxifen injections at P0 and P1. **(B)** Quantification of total EdU-positive cardiomyocytes following EdU administration outlined in Figure 4G represented as a percent of total cardiomyocytes.

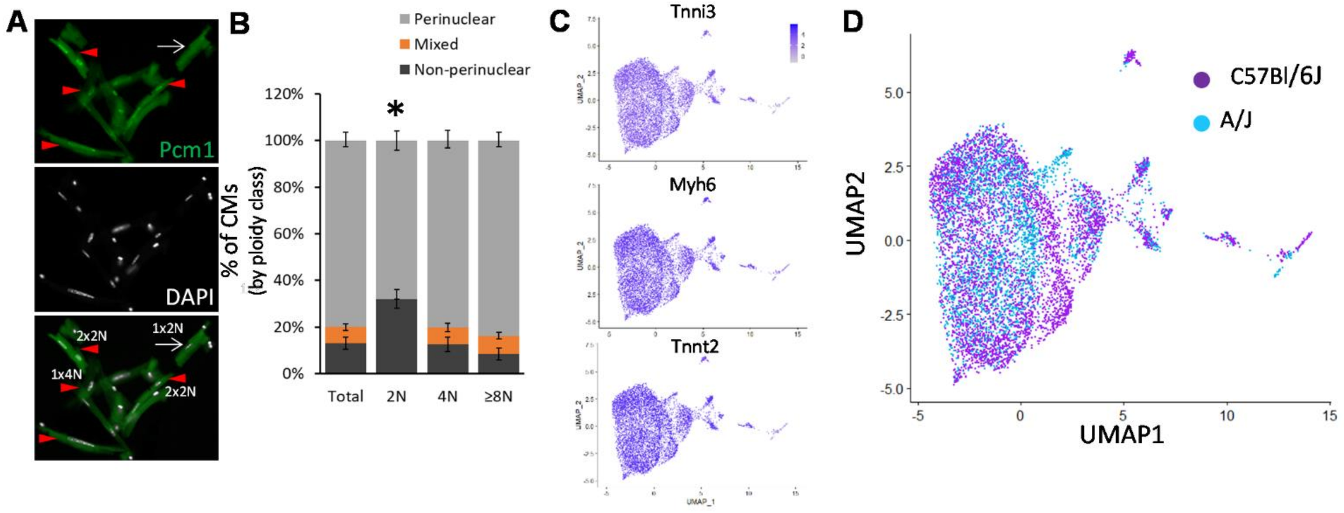


**Supplement to Figure 5. (A)** Immunofluorescent images of Pcm1 (green) and DAPI (greyscale) on single cell suspensions isolated from 6-week A/J hearts. Red arrow heads point to cardiomyocytes with a classic perinuclear staining pattern, while white arrow points to a cardiomyocyte with no perinuclear stain. **(B)** Frequency of observed perinuclear versus non-perinuclear staining pattern for Pcm-1 across ploidy classes. Total = all cardiomyocytes, 2N = 1X2N, 4N = 1X4N + 2X2N, and 8N = all other combinations adding up to 8 or 16N. * indicates P<0.05 for 2N population compared to all others. **(C)** Expression of *Tnni3* (top), *Myh6* (middle), and *Tnnt2* (bottom) across all CM nuclei by single nucleus RNA sequencing. **(D)** UMAP plot distinguishing A/J cardiomyocyte nuclei (blue) from C57Bl/6J nuclei (purple).
